# Genomic analyses of glycine decarboxylase neurogenic mutations, yield a large scale prediction model for prenatal disease

**DOI:** 10.1101/2020.07.28.225292

**Authors:** Joseph D. Farris, Md. Suhail Alam, Arpitha MysoreRajashekara, Kasturi Haldar

## Abstract

Glycine decarboxylase (GLDC) is a mitochondrial protein, hundreds of mutations in which cause a neurometabolic disorder Non-ketotic Hyperglycinemia (NKH), associated with elevation of plasma glycine. But why a mutation induces severe or attenuated neurological disease is poorly understood. We combined a human multiparametric mutation scale that separates severe from attenuated clinical, neurological disease, with new *in silico* tools to assess 238 of 255 NKH mutations in murine GLDC. We unified novel murine and human genome level-analyses across a linear scale of neurological severity, with *in vivo* evidence from mice engineered with a top-ranking attenuated mutation and another mutation >10 times more pathogenic and integrated the data in a model of pre- and post-natal disease outcomes, relevant for over a hundred major and minor neurogenic mutations. Our findings suggest that highly severe neurogenic mutations predict fatal, prenatal disease that can be remedied by metabolic supplementation of dams, in absence of amelioration of persistent and age-dependent elevation of plasma glycine.

## Introduction

Enzymatic defects that alter metabolism may cause complex pathological states including neurological diseases that show a wide range of symptoms such as, seizures, epilepsy, neuropathy, cerebral defects and movement disorders^1-3^. Glycine decarboxylase (GLDC) cleaves glycine in the first step of the mitochondrial glycine cleavage system (GCS). Glycine cleavage yields the formation of 5,10-methylene THF^4^, a key intermediate in folate biosynthesis and one carbon metabolism which is utilized to synthesize nucleotides and proteins and thus broadly impacts cellular homeostasis. Known perturbations of glycine cleavage are associated with cancers, pluripotency, and host susceptibility to viral infection^5-7^. They are also involved in neural tube defects (NTDs) and the rare disorder non-ketotic hyperglycinemia (NKH)^8-11^. The hundreds of mutations in GLDC that give rise to NKH disease, present a unique system to investigate how these mutations underlie NKH, as well as provide models for broader GLDC functions at the cellular and organismal levels.

The incidence of NKH is estimated at 1 in 76,000 births^11^, although higher rates may prevail in remote populations due to founder mutations and a relative lack of genetic diversity^12,13,14^. Loss of GLDC function is the major cause of NKH. This is reflected by an increase of glycine in plasma and cerebral spinal fluid (CSF)^15^. NKH is typically characterized as either severe or attenuated^15,16^. But as summarized in Farris et al. 2020^17^ and references within^15,16^ these disease states are associated with a myriad of over 50 symptoms. Severe NKH induces cerebral defects, intractable seizures, failure to reach developmental milestones and premature death. Elevation of plasma glycine is not predictive of clinical neurological severity, but why this is the case is poorly understood. Confounding factors include blood glycine levels that can be affected by diet. Many patients receive sodium benzoate to reduce circulating glycine levels and this too may affect the perceived lack of correlation to clinical disease. Sodium benzoate is used to control glycine elevation, which is thought to be linked to seizures due to the action of glycine on the NMDA receptor. Dextromethorphan, which antagonizes this receptor, is used to reduce seizure activity. But both sodium benzoate and dextromethorphan are needed at very high doses, leading to concerns of side effects, and they only confer partial benefit if administered early to attenuated patients^18^.

Mouse models help to understand the biology of underlying disease severity. Unfortunately, although there are as many as 255 NKH causing missense mutations in human GLDC, there is a lack of mutation-based mouse models. The large size of the protein and presence of hundreds of mutations also pose difficulties in the optimal selection of mutations likely to induce attenuated or severe disease in a mouse. We previously reported on the development of a mutation-based multiparametric score for human GLDC to distinguish between severe and attenuated clinical, neurological disease^17^. Here, we assemble new *in silico* tools for mapping human mutations to murine GLDC, develop novel, algorithmic approaches to guide selection and validation of neuro-pathogenic mutations engineered in mice and make innovative prediction for hundreds of NKH mutations in prenatal neurological disease.

## Methods

### Homology modeling of GLDC

Homology models were generated for GLDC using the SWISS-model (Swiss Institute of Bioinformatics), which generates homology models as described previously^19-21^. Mouse GLDC was modeled using the crystal structure of P-protein holoenzyme of *Synechocystis* sp. PCC 6803^22^ (PDB ID = 4lhc; sequence identity with mouse = 56.5%) as the template. Human GLDC was modeled as previously described^17^. 242 NKH mutations in human GLDC were mapped onto the mouse GLDC sequence using Clustal Omega sequence alignment^23^ (S Table 1).

### Protein Imaging

3D protein images were generated using the free protein-modeling software Jmol and UCSF Chimera, developed by the Resource for Biocomputing, Visualization, and Informatics at the University of California, San Francisco, with support from NIH P41-GM103311^24^.

### NKH Mutation Analysis

A comprehensive list of NKH-causing missense mutations was compiled through a literature search of previously published mutations and mining of missense mutations catalogued in the ClinVar database hosted by the National Center for Biotechnology Information (NCBI), as described in Farris et al. 2020^17^. Secondary structure location of missense mutations was determined using jMol.

### Murine Multi-Parametric Mutation Score (MMS) for Mouse Mutations

We have previously described the criteria for multiparametric-based mutation assessment of each human GLDC mutation^17^. While the same criteria are relevant for mutations in mouse GLDC, different MMS values were computed and applied, because 8% of residues are not conserved between mouse and human GLDC, and this influences the local environment of amino acids and results in a range of differences in secondary and tertiary structure, which in turn impact ΔΔG values of both stabilizing and destabilizing mutations, as well as helix and sheet mutations and mutations in the dimerization interface. Further, differences arise in values of evolutionary conservation, because the Consurf-generated scores were generated using mouse protein instead of human protein. Finally, there were changes in tRNA because mice have differences in their available tRNAs and codon usage compared to humans (https://doi.org/10.1093/nar/gkv1309). In addition, it was necessary to develop and utilize a novel, Python algorithm which puts mutations through an automated, iterative series of conditional statements to assign scores for 242 murine mutations (S Table 2).

### Weighted Murine Multiparametric Mutation Score (mWMMS) for Mouse Mutations

The 18 MMS parameters were trained on previously described homozygous NKH patient mutations with associated clinical outcome scores (COS)^17^ and homozygous control polymorphisms from the Exome Aggregation Consortium (ExAC) database/gnomAD browser, hosted by the Broad Institute using Python module scikit-learn’s LinearRegression object as previously described by Farris 2020^17^). This yielded the weighted coefficients for all of the 18 parameters (S Table 3). The summation of these weighted parameters yielded a multiparametric mutation score (WMMS) for all 238 of the NKH mutations mapped to mouse GLDC (S Table 1).

### MMS Scoring for all theoretical human and murine missense polymorphisms

MMS comparisons of missense polymorphisms to all theoretical polymorphisms are needed to develop predictive algorithms. A new, custom-designed Python algorithm was used to substitute every nucleotide (A, T, G, or C) at every position of either human *GLDC* or mouse *Gldc* cDNA. Biopython Seq objects were used to handle GLDC cDNA sequence, and nucleotides A, T, G, and C were iteratively substituted at every position. For every substitution, the cDNA was then translated to protein using the Seq.translate() function. Nucleotide substitutions leading to silent mutations or premature stop codons were not analyzed. Nucleotide substitutions leading to protein missense mutations in the first 35 residues of glycine decarboxylase were not analyzed, as positions 1-35 constitute the mitochondrial leader sequence. All other substitutions leading to missense mutations were taken for further analyses. Human and mouse missense mutations and their corresponding cDNA mutations are listed in S Tables 4 and 5. These mutations were scored across the 18 MMS criteria in Python. If a parameter met its condition as described above, it was given a score of 1, except for the conserved substitution parameter, where it was given a score of -1. If the parameter did not meet the condition, it was given a score of 0 (S Table 3). Summation across the 18 parameters in Supplementary tables 4 and 5 yielded an MMS for all human and murine theoretical mutations (S Tables 6 & 7). In these Tables, the minimum score was set to 0. If two nucleotide substitutions led to the same protein mutation with the same MMS score, one was removed so that the same mutation was not included twice in subsequent analyses.

### WMMS for all Theoretical Human and Murine Missense Polymorphisms

WMMS comparisons of missense polymorphisms to all theoretical polymorphisms are needed to develop predictive algorithms. To generate WMMS for all theoretical human and murine missense polymorphisms, human WMMS coefficients (Farris et al. 2020)^17^ were applied a new Python scoring algorithm to all theoretical human mutations. This algorithm contained three parts, i) the cDNA mutation tool described in the prior section which made every possible single nucleotide substitution and yielded their corresponding protein mutation (if any); ii) the MMS-assigning tool described in the prior section which assigned MMS parameter scores for each protein mutation; iii) and the Scikit learn linear regression tool which calculated the weighted parameter coefficients using the homozygous data as described above and applied these weighted coefficients to the MMS parameter values. Summation across parameter values yielded WMMS for all theoretical human polymorphisms (S Table 6). Mouse WMMS coefficients (S Table 3) were applied via the same Python scoring algorithm but modified for mouse protein to all theoretical polymorphisms. Summation across all parameter coefficients yielded WMMS’s for all theoretical mouse polymorphisms (S Table 7).

### Models for ranked prediction for mouse mutations

#### Mutations aligned with attenuated disease

Mouse mutations that correspond to prevalent human mutations associated with attenuated disease^17^ were ranked based on their mWMMS (Score A). The highest mWWMS received a score of 1, the lowest received a score of 0. All other mutations were scored relative to the top-scoring mutation. Mouse mutations were also scored for similarities between their mWMMS and corresponding human mutation WMMS (Score B). No difference was scored as 1 and maximal difference was scored a 0. Mutations were then ranked according to which had the highest sum of Score A and Score B.

#### Parameter prediction for highly pathogenic mutations expected to induce severe neurological disease

Mutations with the top WMMS in the active site region located in the C terminus of the protein (that carries the catalytic Lys759 residue) were assessed for their pathogenic potential based on unique characteristics, in addition to the mWMMS scores. Based on evolutionary conserved functions and mWMMS, S562F, selected as the lead mutation, was used to set the threshold mWMMS of 9.94. Mouse mutations with mWMMS greater than or equal to 9.94, were scored as 1 and those below the threshold were scored 0. Because proline substitutions are highly destabilizing^29^ and could lead to lethality, mutations without proline were scored 1 and those with a proline scored 0. Mouse mutations were also scored for the similarity between their mWMMS and corresponding human mutation WMMS. No difference was scored as 1. Difference between mWMMS and hWMMS was scored 0. Mutations that were located in the active site were scored 1, while those outsides scored 0. Finally, mutations where the mutation was possible in the mouse through single nucleotide substitution were score 1, while those that were not were score 0. The sum of scores of these five parameters was used to calculate the number of mutations with high mWMMS likely to induce severe mouse models of disease.

#### *Gldc* knock-in mutations

CRISPR-Cas9 gene editing technology was used to introduce (i) a moderately pathogenic (A394V, referred to as *Gldc*^*ATT/ATT*^) mutation and (ii) highly pathogenic (G561R, S562F referred to as *Gldc*^*SEV/SEV*^) mutations in the *Gldc* gene. *Gldc*^*ATT/ATT*^ and *Gldc*^*SEV/SEV*^ mutations were custom engineered in the C57BL/6 mouse strain by Jackson Labs (Bar Harbor, ME, USA) and Taconic Biosciences (Albany, NY, USA) respectively. Founder mice were screened for the desired mutations by Sanger sequencing. Germline mutation was confirmed by crossing founders with wild type mice. Homozygous mutant mice were born from heterozygous intercross breeding.

For *Gldc*^*ATT/ATT*^, GCC was mutated to GTG for A394V mutation in the exon 9, creating a restriction enzyme site (GTGAAC) for Hpy166II in the mutant allele. Genotyping of litters were done by PCR amplifying a 753 bp DNA fragment of *Gldc* flanking the mutation site using specific primers (forward: 5’-GTTGCATTTCCGTTTCTGGCT-3’ and reverse: 5’-ACTGCCCTCTTACTTGACCATT-3’). The PCR amplified product was digested with Hpy166II and fragments were resolved by agarose gel electrophoresis. For *Gldc*^*SEV/SEV*,^ G561R, and S562F, were engineered in exon 14 of *Gldc*. Two silent nucleotide substitutions (TGCACC in wild type changed to TGTACA in the mutant), were introduced downstream of the knock-in site, to create a restriction site for BsrG1. For genotyping, a 728 bp DNA fragment of *Gldc* was PCR amplified using primers, forward: 5’-TGCTGTGCTGGGGAGAATTT-3’ and reverse: 5’-TGAACACAGCTACACTCAGCTT-3’ followed by restriction digestion with BsrG1 and analyses by gel electrophoresis.

#### Mouse breeding and other studies

All experiments with mice were performed in compliance with the Institutional Animal Care and Use Committee of the University of Notre Dame.

#### Breeding

Tail biopsies collected on the first day of live and dead births and genotyped. Based on Mendelian inheritance, the expected frequency of wild type, heterozygous and homozygous mutant progenies was set at 25%, 50%, and 25% respectively. Genotype distribution was calculated by dividing the total number of progeny of a genotype by the total progeny in the colony. The Townes mouse model of sickle cell anemia carrying a point mutation (E6V) in the human beta-globin gene and the *Npc1*^*nmf164*^ mouse model of Niemann-Pick type C disease were also bred in-house. Both show autosomal recessive inheritance.

#### Formate supplementation

6-8 weeks old heterozygous mutant female and male were paired. After 48h of pairing, females were separated and sodium formate (30mg/ml) was introduced into the drinking water. The formate solution was replaced weekly. Mice were housed and fed under the standard conditions with a 12 h light/dark cycle. Formate supplementation was stopped after the litter was born.

#### Hydrocephalus

An enlarged cranium with a characteristic dome-shaped head generally accompanied with ataxia and/or reduced mobility was indicative of hydrocephalus, which was later confirmed by the isolation of the brain. Once hydrocephaly was seen, animals were euthanized (there is no effective treatment for hydrocephalus in the mice). Observational studies established that hydrocephalus was seen within the first sixty days of life of mutant mice (at or before the young adult stage).

#### Glycine analyses by mass spectrometry

Blood was collected by cheek bleed from 1-2 month-old *Gldc*^*ATT/ATT*^ mice and age and gender-matched in-colony wild type mice. For *Gldc*^*SEV/SEV*^ mice, blood was collected from 30-60 days and 150-180 day aged *Gldc*^*SEV/SEV*^ and in-colony wild type mice. Plasma was isolated by centrifugation, stored at -80 °C and transported to Wistar Institute Proteomics and Metabolomics Shared Resource for analyses. 20µl of mouse plasma was subjected to cold extraction using 80% MeOH, 20% water, and heavy-labeled internal standards. Samples were analyzed by LC-MS/MS on a Q Exactive HF-X mass spectrometer coupled to a ThermoScientific Vanquish LC System. Metabolites were separated by liquid chromatography and peak areas were extracted using ThermoScientfic Compound Discoverer 3.2 and were normalized to QC sample run. Metabolites were identified from a provided mass list and by MS/MS fragmentation of each metabolite follow by searching the mzCloud database (www.mzcloud.org).

## Results

### Large-scale mapping of human NKH mutations to mouse GLDC

Comparative structural analysis provides a path to understanding functions of conserved proteins. We therefore generated a high-confidence homology model for mouse GLDC protein using *Synechocystis sp. 6803* PLP-bound GLDC (PDB: 4LHC) as a template (Fig. 1A-B). The mouse model received a global mean quality estimate (GMQE) of 0.77 (range = 0 - 1), which compares well to the human model GMQE of 0.77, suggesting a high-confidence homology model to map human mutations to mouse residues. NCBI Blast revealed that the primary sequence of mouse GLDC shares 92% identity of the human orthologue. In addition, mouse GLDC contains an insertion of five glycines (positions 43-47) that are not present in human *GLDC*. Overall the structural features of GLDC’s enzymatic function namely binding of its cofactor pyridoxal phosphate (PLP) at murine Lys759, active site residues (defined as amino acids within 5 A° of the PLP-bound Lys759 or substrate glycine) and the dimerization interface, are conserved in mouse GLDC (Fig. 1D). Using the primary sequences, we mapped 242 of 255 NKH missense mutations in human GLDC to the mouse protein. Thirteen human NKH mutations L207V, R337Q, I440N, I440T, P509A, K574N, E597K, A773P, R790W, C795S, T846I, G860R, and V905G could not be mapped onto mouse GLDC because the affected residue was not conserved and these residues were therefore not further analyzed.

**Figure 1.**
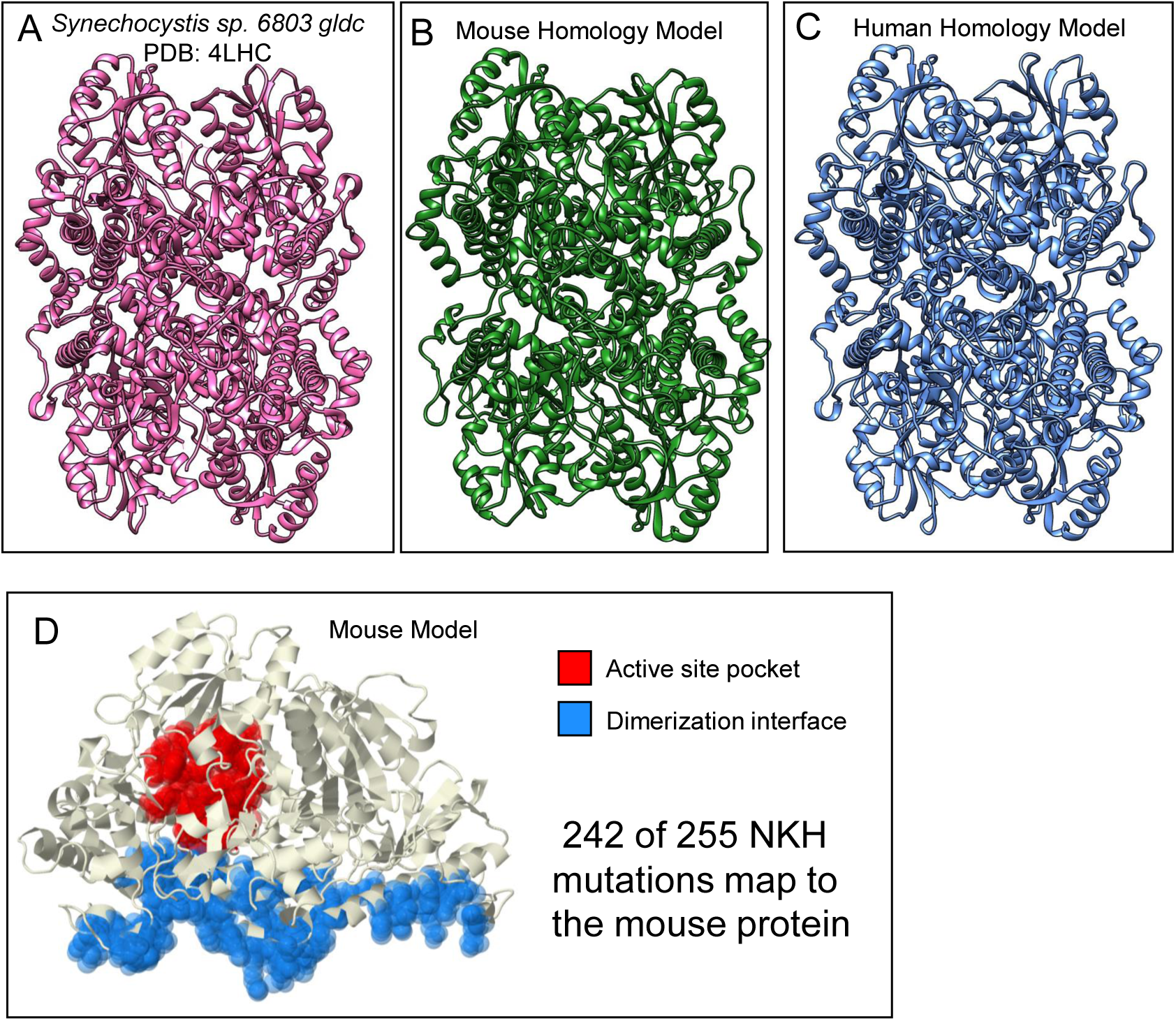
Murine homology model of GLDC-protein. A. A crystal structure of *Synechocystis* sp. PCC 6803 Gldc holoenzyme (PDB = 4LHC) (LHS) was the template used in SWISS-Model to generate (B) the mouse homology model. The mouse model received a global mean quality estimate (GMQE) of 0.77 (range = 0 - 1), showing that this is a high-confidence homology model. This GMQE compares favorably to (C) the human *-* GLDC homology model, which received a GMQE of 0.77. D. Evolution derived functional regions of the mouse GLDC with the active site pocket (red) defined as residues within 5 Angstroms of the PLP-containing active lysine (K759) and the dimerization domain (blue). 242 of 255 NKH mutations map to the mouse protein. The 13 human NKH mutations that could not be mapped onto mouse GLDC were L207V, R337Q, I440N, I440T, P509A, K574N, E597K, A773P, R790W, C795S, T846I, G860R, and V905G.

### Murine Multiparametric Mutation Score

As summarized in Fig. 2A, ΔΔG an important parameter for MMS was assessed by CUPSAT. The SWISS-Model generated mouse GLDC homology model as the input. We defined destabilizing mutations as those with predicted ΔΔG < -1.5 kcal/mol (see Methods). Four of the mutations could not be assessed because they fall either in a short, unstructured region at the N-terminus (D36H, S37N, and G48W) or an unstructured region at the end of the C-terminus (C1007W). But ΔΔG values were obtained for the remaining 238 mutations (see S Table 1). 53 mutations were predicted to destabilize GLDC. 48 were predicted to be very destabilizing (< -5 kcal/mol). 50 mutations were predicted to be stabilizing, and the remaining 87 mutations were predicted to have a negligible effect on protein stability. However, MMS and WMMS, rather than ΔΔG alone were needed to separate severe and attenuated human neurogenic mutations^17^. We therefore computed MMS values for the 242 murine mutations (Fig. 2C; S Table 2). The distribution of murine MMS scores in the murine GLDC protein is shown in Fig. 2D. Only 4 of 242 mutations received a score of 0. Mutations with scores of 1-2 were considered mild (N = 51), 3-4, moderate (N = 75), and ≥5 severe (N=112) (S Table 1). These are consistent with prior findings reported for human GLDC^17^ but also reflect important differences in the mouse protein. In summary, environmental differences at the level of local amino acids in secondary and tertiary structures, as well as murine-specific effects that arise due to changes in evolutionary and protein translation models, may alter the effects of a significant fraction of mutations (∼35%) between human and mouse proteins.

**Figure 2.**
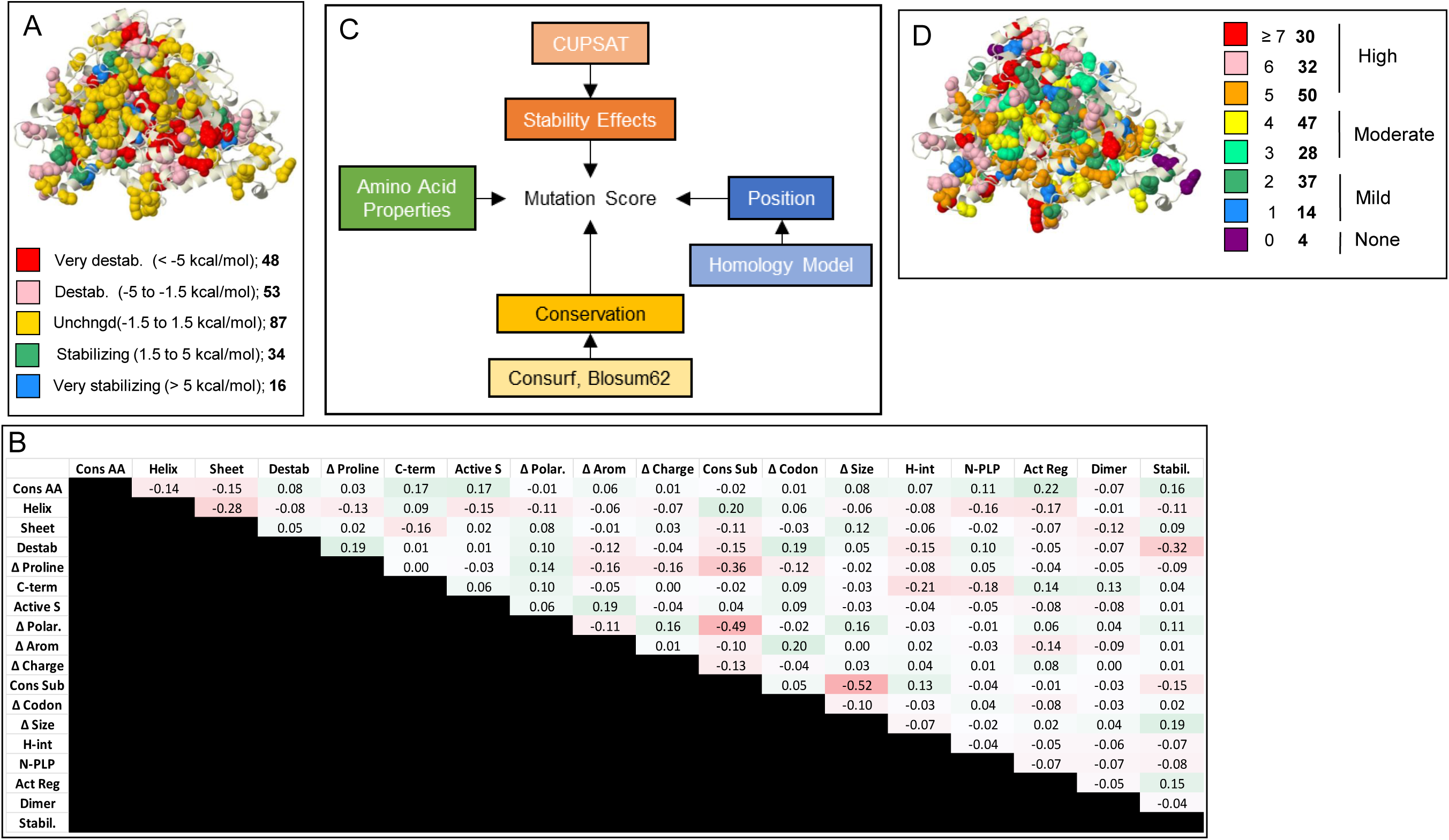
Murine Multiparametric Mutation Score (MMS). [A] Stability effects. CUPSAT-predicted stability effect of NKH mutations on GLDC-protein assessed by ΔΔG for mouse GLDC. For very destabilizing and destabilizing mutations, predicted ΔΔG were <-5.0 and -5.0 to -1.5 kcal/mol respectively. Predicted ΔΔG predicted to be unchanged, stabilizing or very stabilizing are as shown. Number of mutations in each ΔΔG category shown in bold. [B] Phi correlations of 18 parameters predicted to impact protein stability. Most parameters have no correlation with a phi-value of < 0.3. Change in proline, size, and polarity have weak negative correlations with the conservation of amino acid change parameter, indicating that alterations in these amino acid properties are not well-tolerated through evolution. [C] Design for calculating MMS. In total, 18 parameters were used from 4 general categories of stability effects, mutation position, conservation, and change in amino acid properties (see Methods). [D] MMS was calculated for mouse GLDC from a summation of the 18 parameters, each with a weight of 1 except conservation of substitution, which was weighted -1. 3-dimensional distribution of MMS’s shown for all known NKH missense mutations. Key indicates the number of amino acids associated with each MMS range (shown in bold).

### Mouse Weighted Multiparametric Mutation Scores (mWMMS) to guide the design of preclinical mouse models

In mouse studies, we expected to apply mutation scores to develop models where both alleles would be mutated to generate homozygous mutations. We therefore used previously described clinical neurological disease scores associated with homozygous human mutations ^17^ to generate appropriate weighting parameters (S Table 3) and applied these to 238 mutations that received an MMS score. The resulting mWMMS provides an assessment of neurogenicity for 238 mouse mutations, that ranged from -7.8 to 14.4 as presented in S Table 1. Higher mWMMS predict increased neurogenicity while those below zero predict reduced pathogenesis. Nonetheless, a significant portion of mWMMS are different from hWMMS, and determining which of (hundreds of) ordered mutations will successfully induce neurological disease when engineered into an animal remains a challenge. Not all human pathogenic mutations maybe efficiently expressed in a mouse, the most severe mutations may result in embryonic lethality and the neurogenicity of disease mutations compared to benign mutations may be different in humans compared to mice.

To additionally improve our understanding of the relationships between human and mouse proteins and their neurogenic mutations, we developed a new algorithmic pipeline which introduces mutations at the DNA level, finds corresponding protein mutations, generates an MMS for each protein mutation, and weights the MMS to generate the WMMS (see Methods). We utilized this pipeline to further explore the WMMS in both human and mouse proteins by analyzing (i) all theoretical mutations possible in full length proteins (S Tables 4-7) (ii) known neurogenic mutations alone (S Table 1) and (iii) pathogenic and benign mutations (where benign was defined as the subset yielded when pathogenic mutations were subtracted from all theoretical mutations). Our analyses reveal exact correspondence of WMMS in 67.3% of human and mouse mutations throughout the GLDC protein (S Table 7, Fig 3A) as well as in 65.3% of neurogenic human mutations and corresponding counterparts in mice (S Table 1, Fig. 3B). Together the findings suggest that despite 92% identity between human and mouse GLDC, only 60-70% of human NKH mutations may indeed be well mimicked in mice.

**Figure 3.**
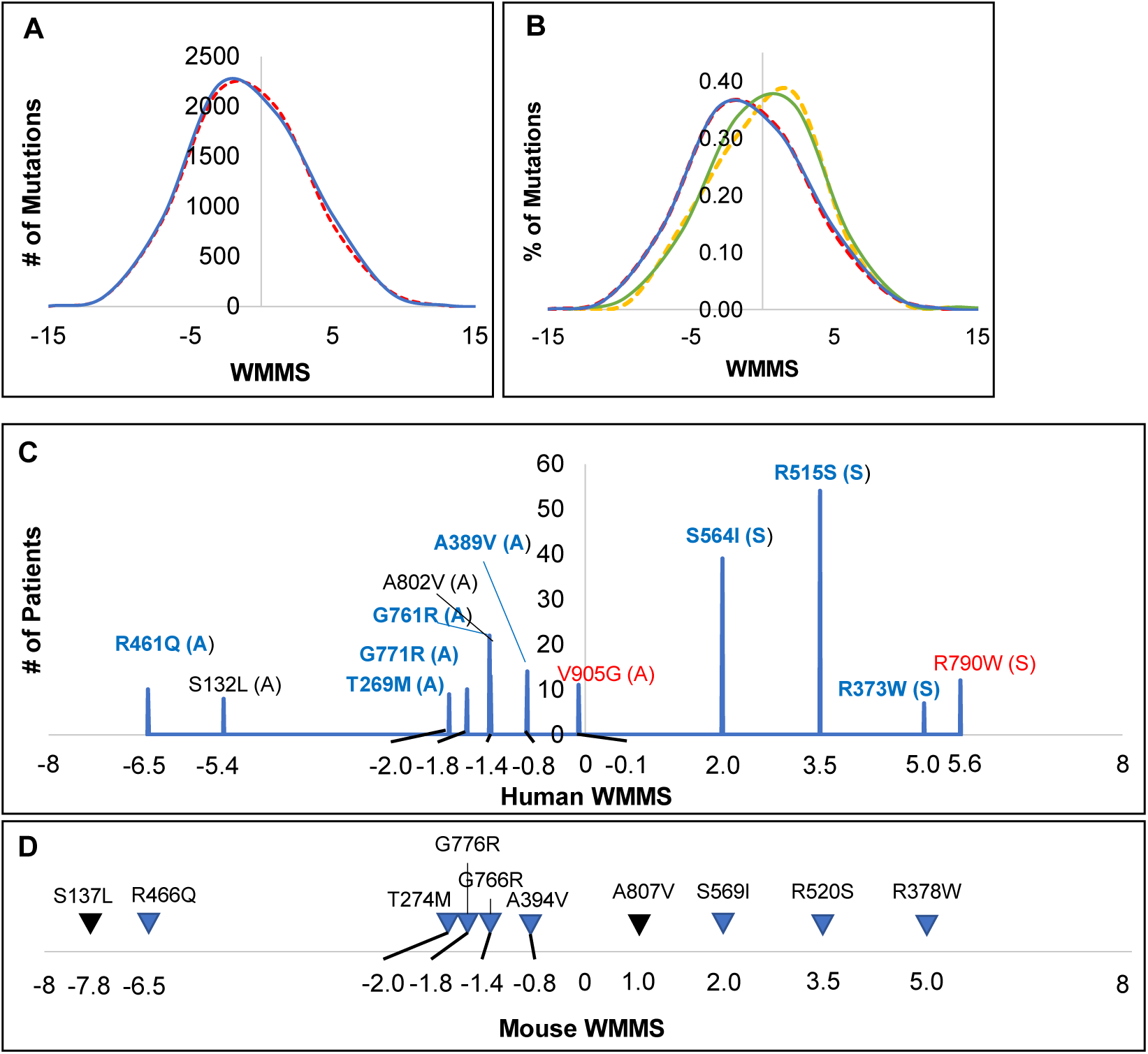
Comparative analyses of Weighted Multiparametric Mutation Scores (WWMS) of Mouse and Human Polymorphisms. **[A]** The distribution of all theoretical missense mutations caused by single-nucleotide substitutions in human *GLDC* cDNA (red dashed line) and mouse *Gldc* cDNA (blue solid line). **[B]** WMMS comparison of known pathogenic mutations in human (yellow dashed lines) and their correlates in mouse (green solid line) vs theoretical mutations whose pathogenicity is unknown in both human (red dashed line) and mouse (blue solid line) glycine decarboxylase. **[C]** The twelve most prevalent human mutations and their hWMMS scores. Attenuated (“A”) or severe (“S”) predictions were assigned based on the mutation’s predicted clinical outcome score (“A” <5; “S” ≥ 5) based on Farris et al. 2020^17^. Mutations for which the human residue is not conserved in mouse GLDC are shown in red. **[D]** Mouse counterparts for the most prevalent human mutations and their mWMMS scores. Mutations for which mWMMS is different from hWMMS are shown in black.

Since prevalent mutations are best characterized and understood, we used them to prioritize mutant selection to develop preclinical models. We therefore examined the WMMS of the 12 most prevalent human NKH mutations^17^ and their counterparts in the mouse (Fig. 3C; S Table 8-9). As expected, WMMS scores of the four human neurogenic mutations predicted to be severe were higher than the eight mutations of predicted attenuated disease (Fig. 3C). One mutation (R790W) associated with severe and another (V905G) associated with attenuated clinical disease were not conserved in mice. Of the remaining ten, the hWMMS scores of eight were identical to that of their mouse counterparts (Fig. 3D). The human mutation A802V showed a hWMMS of -0.8, which increased to mWMMS of 1.0 for its corresponding mouse mutation A807V. A second human mutation S132L showed hWMMS of -5.4 that decreased to mWMMS of -7.8 in its mouse counterpart S137L. Differences between hWMMS and mWMMS (which arise due to intrinsic differences in their respective MMS) suggested that parameters of these human mutations were not optimally reflected in the mouse.

Our goal was to utilize the mWMMS to maximize the potential for a mouse mutation to induce neurological disease that can be productively studied for severity and disease correlates over several weeks. We therefore reasoned that a robust path to achieve the first mouse with a benchmark model of disease would be to select a mutation with the highest mWMMS aligned with attenuated human disease (Fig. 4A). Of the 7 prevalent attenuated human mutations and their mouse correlates shown in Fig. 4B, mouse A807V reveals the highest mWMMS of 0.96 and S137L yields the lowest mWMMS of -7.77. However, after incorporating differences between mWMMS and hWMMS (compared in Fig. 4C), A394V received a ranked prediction of 1, with A807V and S137L ranked 6 and 7 respectively (Fig. 4D). Therefore, A394V mWMMS -0.87 was the topmost hit of the prediction model and progressed to *in vivo* validation

**Figure 4.**
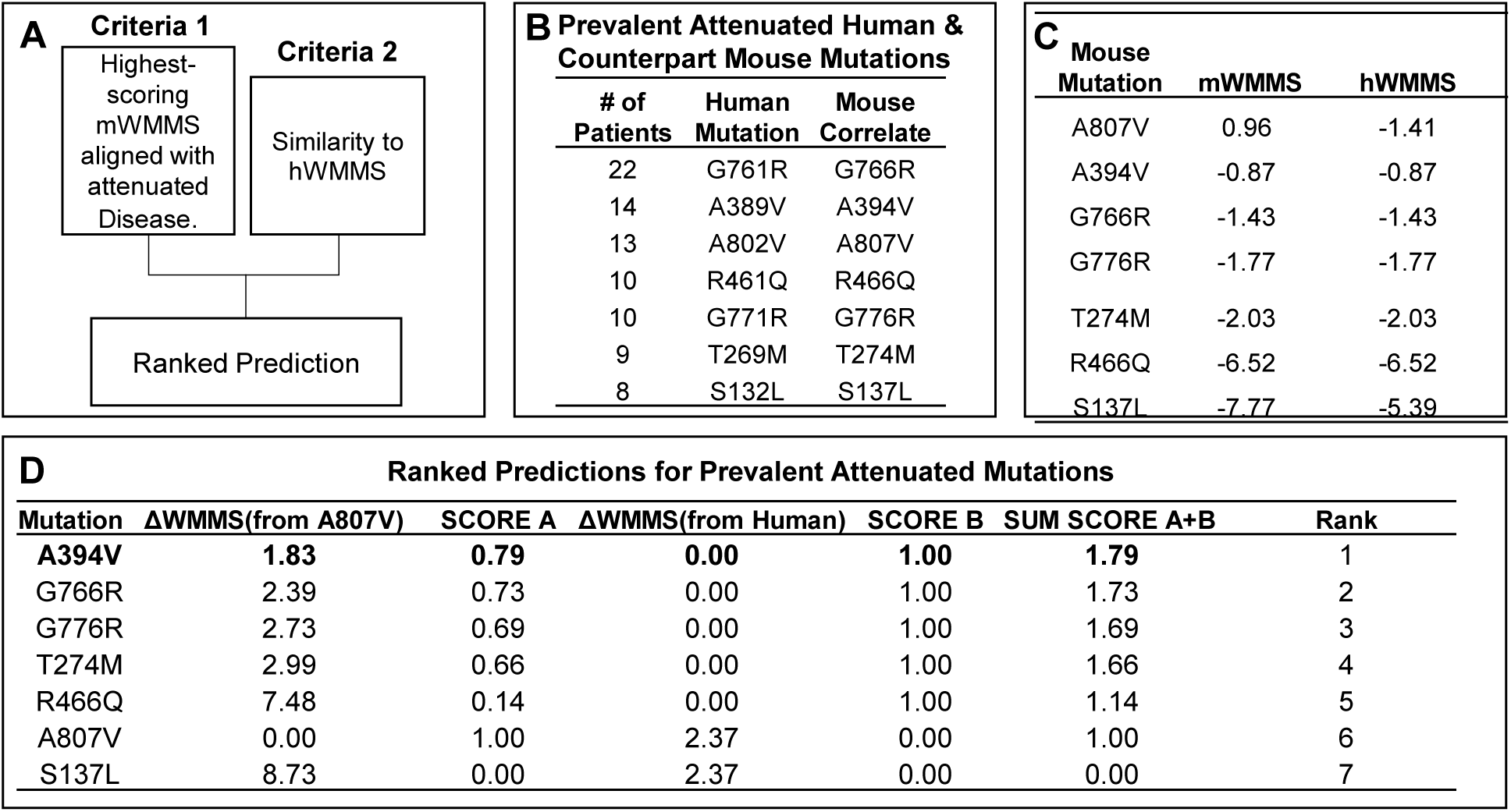
A predictive model to rank mouse mutations aligned with attenuated disease for validation studies. **[A]** WMMS (mouse, and human, hWMMS) ranking criteria for inclusion in predictive model. **[B]** Top 7 most prevalent attenuated mutations in NKH patients and correspondence position in mouse *Gldc*. **[C]** Mouse mutation, corresponding mWMMS and hWMMS of human counterparts **[D]** Predictive Ranking. SCORE A was assigned based difference in WMMS from the most severe attenuated mutation A807V. SCORE B was assigned based on the difference in WMMS between the mouse and human scores. Ranks were assigned based on the highest sum of SCORE A and SCORE B leading to the selection of A394V to be forwarded for *in vivo* validation.

### Validation of A394V as a mutation that induces elevation of glycine and severe cerebral defect in an attenuated mouse model of NKH

A394V was genetically engineered using CRISPR-Cas9 in C57BL/6J mice. For detailed procedures, please see Methods. The genomic location and the strategy for genotyping colonies are summarized in Fig. 5A. Alanine 394 in exon 9 was mutated to Valine (A394V) using two nucleotide substitutions that enabled Hpy16II restriction analysis via characteristic band patterns for wild type heterozygotes and homozygous mutants. Heterozygotes were crossed to yield homozygous A394V mutants, which were designated *Gldc*^*ATT/ATT*^. As shown in Fig. 5B, out of a total of 87 progeny, 17 (19.5%) were *Gldc*^+/+^, 55 (62%) were *Gldc*^*ATT/+*^ and 16 (18.4%) were *Gldc*^*ATT/ATT*^. The Chi-square test suggested no significant differences (χ^2^>0.05) between the observed and expected numbers of animals of each genotype (Fig. 5C). There was no significant gender bias within each genotype (χ^2^>0.05; Fig. 5C). Mouse colonies of two other monogenetic autosomal recessive disorders, sickle cell anemia (carrying a point mutation in the β -globin gene;^30^ and Niemann-Pick Type C disease (carrying a point mutation in *Npc1* gene ^31,32^) yielded homozygous mutant progeny at rates between 21-22% (Fig. 5B). While yield of *Gldc*^*ATT/ATT*^ at 18.4% is measurably lower, the strain could nonetheless be productively maintained and studied, and was therefore further characterized.

**Figure 5.**
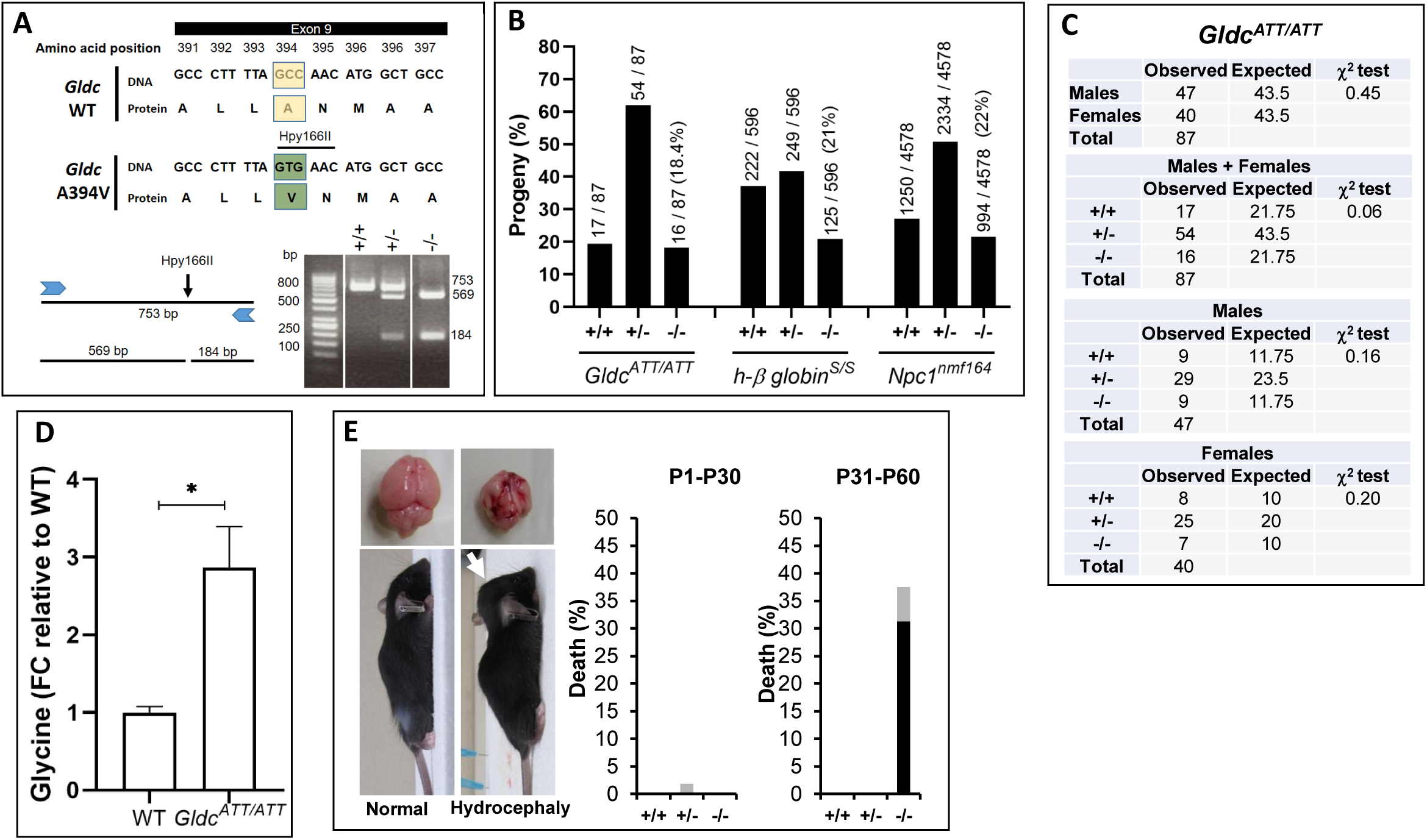
Validation of A394V mutation in an attenuated mouse model of NKH. [**A**] A394V was engineered in exon 9 of *Gldc*. A Hpy166II restriction enzyme site (GTGAAC), generated while engineering was utilized for genotyping (Methods). Wild type (+/+), heterozygous (+/-) and homozygous (-/-) mutants were respectively recognized by PCR followed by restriction digestion yielding one fragment of 753 bp, three fragments of 753, 569, and 184 bp and two fragments of 569 and 184 bp. **[B]** Percent yield of mouse progeny from mating of heterozygous *Gldc*^*ATT/+*^ pairs. The fraction indicates the number in each genotype divided by the total number in the colony. Percent yield of progeny from mouse models of sickle cell anemia (human β-globin^s/s^) and Niemann-Pick type C disease (*Npc1*^*nmf164*^). are shown as reference. [**C**] Gender distribution in *Gldc*^*ATT/ATT*^. The expected numbers (25% wt, 50% heterozygotes and 25% homozygous) are based on Mendelian inheritance. **[D]** Fold difference in plasma glycine levels in *Gldc*^*ATT/ATT*^ and wild type (WT) at 30 - 60 day. n=3 (2F, 1M) mice per group. The data is mean±SEM. *p<0.05, unpaired Student’s t-test. **[E]** Hydrocephalus, a characteristic dome-shaped head (arrow) results in brain atrophy in *Gldc*^*ATT/ATT*^ from day 1-30 (P1-30) and day 31-60 (P31-60). Black bars, percent death due to hydrocephalus; grey bars, percent death due to other causes. +/+, wt; +/- heterozygous mutant and -/-, homozygous mutant.

To investigate whether *Gldc*^*ATT/ATT*^ display correlates of NKH disease, we assessed mice for plasma glycine levels, elevation of which is the characteristic defect of the underlying genetic defect. As shown in Fig. 5D, in mice at or below 60 days of age, we detected a 2.9-fold increase in plasma glycine levels in *Gldc*^*ATT/ATT*^ compared to wild type littermates, For neurological disease, we reviewed the wide range of neurological symptoms and brain malformations associated with human NKH (Farris et al. 2020 and references within)^17^. The most readily discernible symptom in *Gldc*^*ATT/ATT*^ was hydrocephaly. We used it because it can be detected as a characteristic enlargement of the head (due brain atrophy and abnormal accumulation of cerebrospinal fluid), without intrusive methods and therefore enabled rapid scoring of a brain defect prior to sacrificing animals. As shown in Fig. 5E, 31% of *Gldc*^*ATT/ATT*^ developed hydrocephaly detectable between 30 to 60 days (at late juvenile to young adult stages), compared to none in heterozygotes or wild type mice. Hydrocephaly was also the principal cause of death. Together these data suggested that A394V mutation induced an attenuated disease model with mild and moderate defects respectively in prenatal and early postnatal neurological development.

### Prediction and selection of a highly severe mutation for validation studies

Validation of A394V suggested that any mutation with mWMMS > -0.87 should display a measurable level of prenatal disease and postnatal neurogenicity. But we wanted to understand how to model the upper range of disease severity to encompass the largest possible number of neurogenic mutations that lie between -0.87 and the highest mWMMS that could be productively analyzed in a mouse.

The top mWMMS for all theoretical mutations is 14.4 (Fig. 3A, S Table 7). But the highest WMMS seen for prevalent severe mutations, was much lower (5.0; Fig. 3C-D). We subsequently examined 18 mutations found as homozygous alleles in patients (for which we could directly assign a clinical outcomes score (COS ranging from 0-12). For 15 mutations, the hWMMS was equal to the mWMMS (Fig. 6A). Eight correspond to clinically attenuated human mutations (COS <5) with hWMMS/mWMMS that range from -2.03 – 1.13. Seven mutations correspond to clinically severe (COS >5) and show hWMMS/mWMMS from 0.37 – 7.44 (Fig. 6A). Except for one outlier (at 0.37), the mutations with severe COS showed a WMMS range higher than attenuated mutations. A best fit plot revealed a linear relationship between WMMS and COS for attenuated and severe neurogenic mutations (Fig. 6B). WMMS of 1.3 defines the transition from attenuated to severe neurological disease based on the COS cutoff of 5. The linear relationship (r^2^ 0.9) between WMMS and COS, also suggested that outcomes of a single severe mutation in mice taken in conjunction with those of the attenuated mutations A394V, would be sufficient to predict outcomes of a wide range of neurogenic mutations. Further, the severe mutation could be of clinical origin or synthetic in nature (derived from all theoretical mutations shown in Fig. 3A). We also reasoned that highly severe mutations (WMMS greater than or equal to 10) were unlikely to be prevalent as homozygous mutations in patients. However, modeling them in mice would help to better understand their properties (that may be difficult to study in humans).

**Figure 6.**
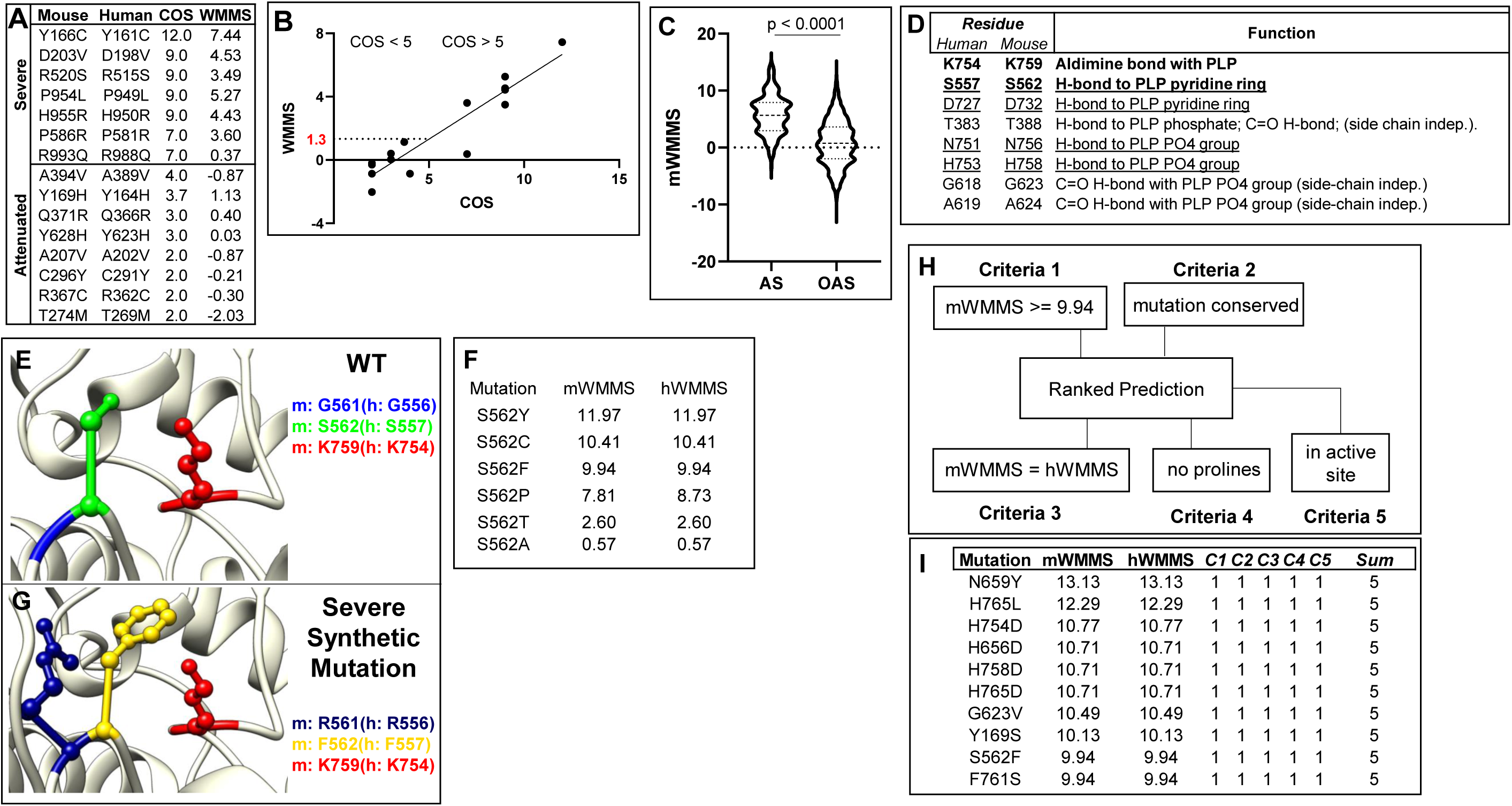
Prediction and selection of a pathogenic mutation to induce a mouse model of severe disease. **[A]** Fifteen homozygous human mutations where hWMMS is the same as mWMMS and their assigned human clinical outcome scores (COS) from Farris et al. 2020^17^ are presented. **[B]** Distribution of WMMS for mutations (from panel A) predicted to be attenuated (COS < 5) and severe (COS ≥ 5). The distribution follows the trend y = 0.76*x - 2.5; r^2^ 0.9. The WMMS transition from severe to attenuated state is predicted to be linear and projected to occur at a score of 1.3. **[C]** Distribution of mWMMS values for theoretical mouse mutations inside (N = 204) and outside (N = 5,243) the GLDC the active site. Active site mutations have a mean mWMMS of 5.8, while mutations outside the active site have a mean mWMMS of 0.6. The two distributions are significantly different (P < 0.0001) by unpaired student’s t-test. **[D]** Active site residues that form hydrogen-bonds relevant for catalysis. Active site lysine and lead severe mutation candidate, S562F, are shown in bold. Side-chain dependent hydrogen-bonds are underlined. **[E]** The position of G561 (blue) and S562 (green) relative to active site K759 (red) **[F]** Possible S562 substitutions, mWMMS and corresponding hWMMS’s are shown for each substitution. **[G]** Representation of the double synthetic mutations G561R (dark blue), S562F (yellow) in GLDC. Active site lysine (K759) is shown in red. **[H]** WMMS (mouse and human) and other molecular ranking criteria for inclusion in severe mutation selection model. **[I]** The 10 theoretical mutations that scored in all five criteria laid out in 6F. Note, S562C and S562Y shown in panel D are excluded because they were previously not predicted to disrupt the hydrogen bond of S562.

Since we could not be guided by highly severe known human severe neurogenic mutations, we resorted to developing a highly severe synthetic mutant. To do so, we began by examining the active site region, since mutations here may closely impact the catalytic activity. As expected, active site region mutations show a higher median mWMMS than mutations not in the active site (Fig. 6C; S Table 10). The active site region contains the PLP cofactor bound to the catalytic residue Lys759. Mutation of Lys759 is likely to be lethal. But as shown in Fig. 6D, based on comparison to *Synechocystis* GLDC crystal structure^22^, the active region also contains a number of residues that stabilize PLP via hydrogen bonding (H-bonding) that we examined because they are likely to be important for catalytic activity. Hydrogen bonding is known to be important for function in a wide range of PLP enzymes^33^. However, H-bonding by T338, G623 and A624 is through the backbone carbonyl group, and therefore is side chain-independent. In the *Synechocystis* GLDC model, S529 makes a hydrogen bond to the PLP pyridine ring^22^. Its mouse counterpart S562 was found to be appropriately positioned with respect to the active site Lys759 in a 3-D model (Fig. 6E). Additional residues such as D732 H-bond to the PLP pyridine ring, while N756 and H758 H-bond to PLP phosphate groups. But mutation in S562 is expected to have the least impact on folding of the active site because it lies furthest linearly from K759 and therefore was preferred. Analyses of synthetic mutations in S562 suggested possible substitutions S562Y, S562C, S562F, S562P, S562T, and S562A (Fig. 6F). S562C, S562Y and S562F had higher mWMMS (>9.94), but of these S562F is expected to most effectively disrupt hydrogen bonding^34^. Because it is a contextual property, H-bonding was not included in the parameter S Table 3 and therefore not included in neither the MMS nor WMMS. But due to its importance for catalytic activity, H-bonding coupled with a high mWMMS (9.94) was used to prioritize S562F over S562Y and S562C and with higher mWMMS (11.97 and 10.41 respectively). The placement of S562F relative to the active site in a 3D model is shown in Fig. 6G.

To benchmark our consideration of S562F as the first severe mutation, we developed a model (Fig. 6H) to compare all theoretical mutations with mWMMS that were (i) greater than or equal to 9.94, (ii) conserved across mouse and human species, (iii) aligned with hWMMS, (iv) lacking prolines (because of the extreme destabilization conferred by this amino acid^29^ which may engender lethality) and (v) located to the active site. The presence of each parameter was scored a 1 and absence was scored 0. As shown in Fig. 6I, S562F was not amongst the highest mWMMS score but it was among 10 mutations that received the top score of 5 and represented the top 0.2% (10 of 5447) of total theoretical mutations (S Table 11).

To retain the attributes of S562F but increase severity, we used an adjacent second mutation (G561R, mWMMS 5.9) to create a double mutation G561R,S562F (Fig. 6G). S562F introduces an aromatic residue in the core of the protein. The introduction of arginine via G561R is expected to be highly antagonizing as a second mutation. However, the double mutation could not be scored and assigned a mWMMS because the utilized parameter weights were trained on GLDC’s with single-point mutations. Nonetheless, based on the scores of each mutation and the H bonding disruptive properties ascribed to S562F, the G561R,S562F was expected to be significantly more pathogenic than WMMS of 10. The major caveat for G561R,S562F was that it might be lethal and therefore possibly unproductive for study, but we reasoned that if that were the case, we could separately test each mutation to establish their expected disease severity. Further, our predictive model (Fig. 6 H-I) suggested multiple additional mutations that may also present robust alternatives to induce severe neuro-pathogenicity.

### Engineering and analyses of a highly severe mutation-based mouse model of NKH

G561R,S562F was genetically engineered using CRISPR-Cas9 in C57BL/6 mice. For detailed procedures, please see Methods. The genomic locus and genotyping strategy are shown in Fig. 7A. Two silent mutations introduced immediately downstream of the mutation site created a restriction site for BsrG1 (TGTACA). Digestion yielded characteristic band patterns for wild type heterozygotes and homozygous mutants. Heterozygote crossing was used to generate homozygous mutants G561R,S562F that were designated *Gldc*^*SEV/SEV*^. A total of 10 *Gldc*^*SEV/SEV*^ pups were born in 404 progeny from 122 litters over 22 months (Fig. 7B). This reflects progeny yields of 2.5%, which is 10-fold lower than the maximum theoretical yield of 25% expected of homozygous mutants. None of the *Gldc*^*SEV/SEV*^ pups survived beyond 24 hours. Since all genotyping samples were collected on postnatal day 1, we attributed the reduced number of *Gldc*^*SEV/SEV*^ mice to severe prenatal and/or very early birth defects.

**Figure 7.**
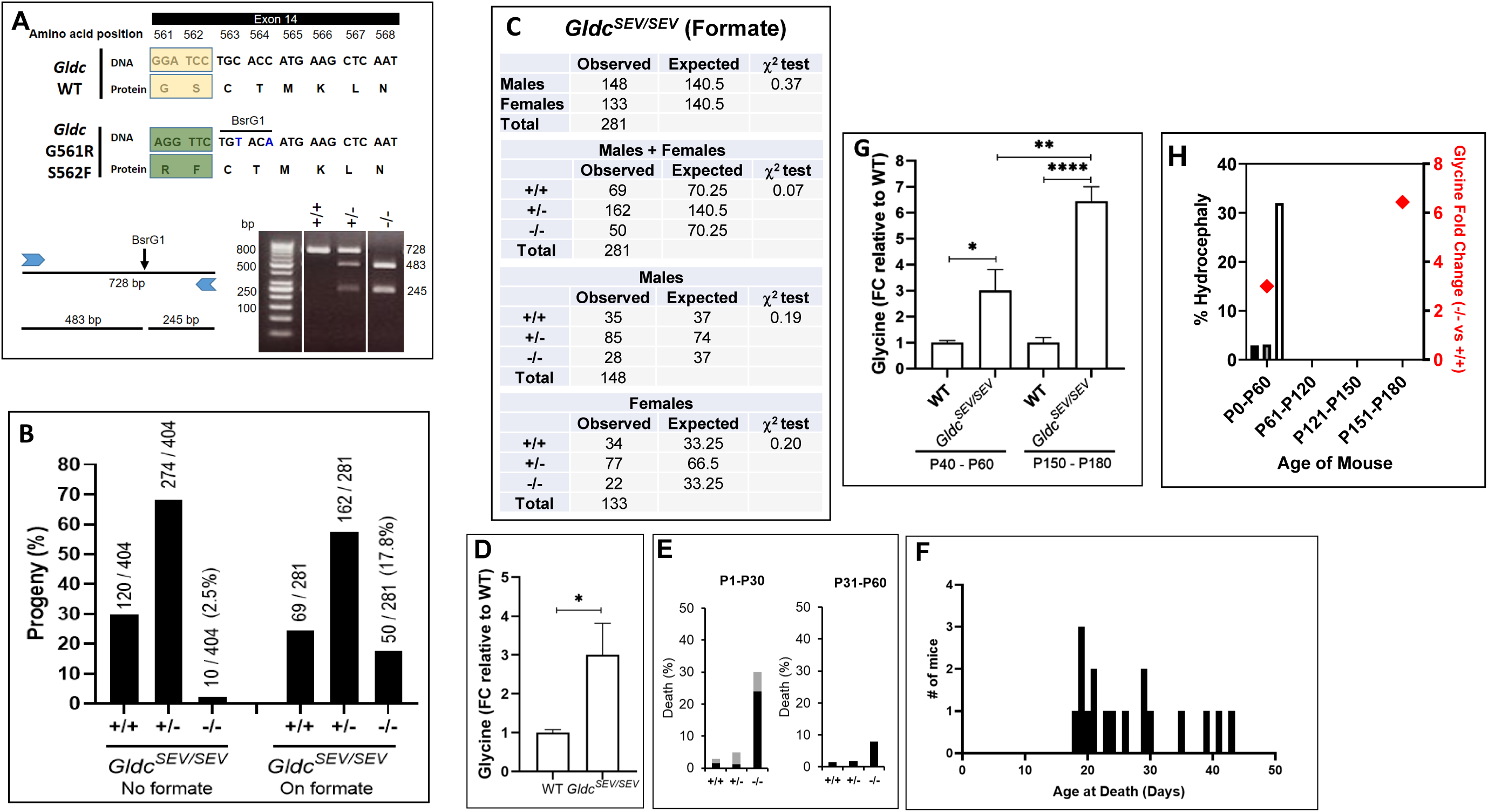
Validation of a highly pathogenic mutation in a severe mouse model of NKH. **[A]** G561R S562F, was engineered in exon 14 of *Gldc*. Two silent nucleotide substitutions (TGCACC in wild type changed to TGTACA in the mutant), were introduced downstream of the knock-in region, to create a BsrG1 restriction site. Wild type (+/+), heterozygous (+/-) and homozygous (-/-) mutants were respectively recognized by PCR amplification followed by restriction digestion yielding one fragment of 728 bp, three fragments of 728, 483, and 245 bp and two fragments of 483 and 245 bp. **[B]** Percent yield of mouse progeny from mating of heterozygous *Gldc*^*SEV/+*^ pairs engineered with G561R,S562F. The fraction indicates the number in each genotype divided by the total number in the colony. Dams were treated with or without formate as shown. **[C]** Gender distribution in *Gldc*^*SEV/SEV*^. The expected numbers (25% wt, 50% heterozygotes and 25% homozygous) are based on Mendelian inheritance. **[D]** Fold difference in plasma glycine levels in *Gldc*^*ATT/ATT*^ and wild type (WT) at 30 - 60 day. n=6 (3F, 3M) mice per group. The data is mean+/-SEM. *p<0.05, unpaired Student’s t-test. **[E]** Hydrocephalus, in *Gldc*^*SEV/SEV*^ from day 1-30 (P1-30) and day 31-60 (P31-60). Black bars, percent death due to hydrocephalus; grey bars, percent death due to other causes. +/+, wt; +/- heterozygous mutant and -/-, homozygous mutant. [**F]**. Hydrocephalus incidence as a function of age. [**G**] Comparative analyses of plasma glycine levels in *Gldc*^*SEV/SEV*^ mice at 40-60 d and 150-180 days. Age-matched wt served as controls. n=6 (3M +3F) in each group. Mean±SEM values are shown. *p<0.05, unpaired Student’s t-test. [**H]** Comparative analyses of the time course of hydrocephalous and glycine elevation from day 0 to 180.

In a gene trap null mouse model, *Gldc* was shown to be required for neural tube development and closing, but this requirement could be circumvented by formate supplementation^35^. We therefore introduced formate in the drinking water of *Gldc*^*SEV/+*^ dams from the outset of pregnancy (Methods). Of the resulting 278 progeny, 50 were *Gldc*^*SEV/SEV*^ (Fig. 7B). Statistical analysis showed no significant difference (χ^2^>0.05) between the observed and expected number of progeny for each genotype and gender (Fig. 7C). Formate supplementation did not adversely affect the birth of *Gldc*^*+/+*^ and *Gldc*^*SEV/+*^ mice since their yields and gender were within the normal range (Fig. 7C). These data strongly support that effect of the double mutation G561R,S562F on embryonic development was due to a depletion in one carbon metabolites, which arises from reduction in glycine cleavage and can be circumvented by supplementing dietary formate of dams.

We next investigated whether *Gldc*^*SEV/SEV*^ progeny display post-natal correlates of NKH disease. Over the first 60 days, plasma glycine levels increased 3-fold in mutants compared to wild type mice (Fig. 7D). Further, 34% of *Gldc*^*SEV/SEV*^ developed hydrocephaly (Fig. 7E). Postnatal correlates of glycine and the fraction of hydrocephaly of *Gldc*^*SEV/SEV*^ treated with formate were similar to that seen in *Gldc*^*ATT/ATT*^. In *Gldc*^*SEV/SEV*^, hydrocephalus and associated death was observed between 15 and 45 days (Fig. 7F). Notably, the *Gldc*^*SEV/SEV*^ mice that developed to a mature age (> 3 months) had substantially increased glycine levels between 150-180 days (compared to 60 days; Fig 7G), suggesting animals matured and aged without fatal cerebral defects in spite of aberrantly higher levels of plasma glycine (Fig. 7H).

### Predictive outcomes for major and minor NKH human disease mutations

We have in this study developed several new computational approaches to establish an *in silico* murine multiparametric mutation scale linearly proportional to disease severity, based on which we engineered two mice, one with the A394V mutation (mWMMS -0.87), another carrying a G561R,S562F double mutation (mWMMS > 10). A394V induced 26% prenatal lethality and 31% hydrocephalus in the postnatal stage. G561R,S562F showed 90% prenatal lethality and no viable pups. When the dams were treated with formate, prenatal lethality reduced to 29% and hydrocephalus was seen in 34% of the mice. These data predict that increase in WMMS is reflected by an increase in fatal prenatal disease, rather than post-natal hydrocephalus. Notably, 34% hydrocephalus seen in the double mutant after exposure to formate, may also reflect prenatal defects that are not corrected by the treatment of dams.

To apply these findings at a larger scale, we examined NKH mutations with WMMS equal to or greater than -0.87 and identical to the corresponding mouse WMMS. As shown in Fig. 8A, 126 mutations were identified and their distribution across the protein suggested they displayed no overt positional bias (see also S Table 12). Two out of 126 mutations received scores above 10 (Fig. 8A, green, encircled in red), but these were due to proline substitutions (and therefore not retained for subsequent analyses). 33 mutations were predicted to be attenuated (WMMS range -0.87 and 1.3) and the remaining 91 were deemed severe (WMMS between 1.3 and 10). 81 of the severe mutations (WMMS 1.3-7.4) are the most relevant to the vast majority of presently known neurogenic missense mutations associated with severe NKH in humans.

**Figure 8.**
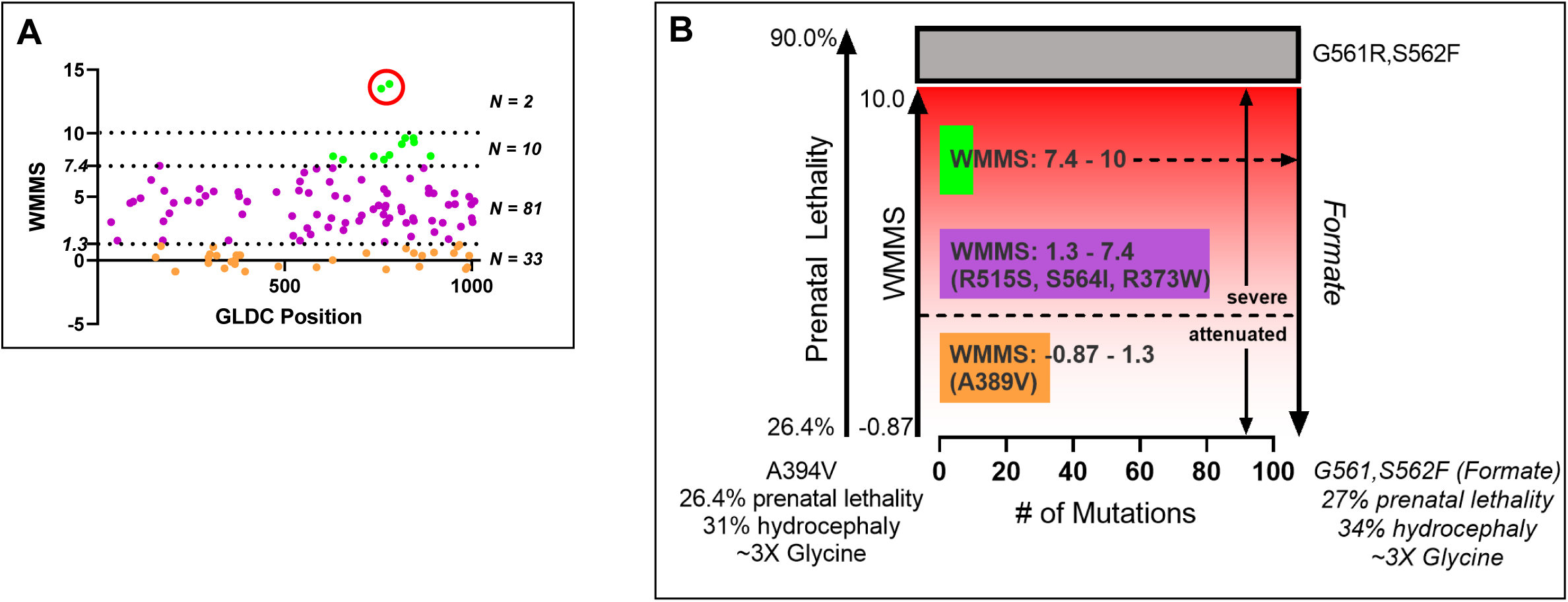
Predictive outcomes for major and minor NKH human neurogenic mutations. [A]. Mouse counterparts of human NKH mutations with identical mWMMS and hWMMS and WMMS is ≥ -0.87 displayed as a function of their distribution across GLDC protein. Mutations ranging in the attenuated range from WMMS -0.87 to 1.3 (N = 33) are shown in orange. Severe mutations range from 1.3 to 7.4 (N = 81) are shown in purple. 10 mutations (shown in green) lie between WMMS 7.4 (the highest WMMS score of the most severe homozygous patient mutation) and 10. The two mutations with scores above 10 (green circled in red) lie beyond the WMMS range of - 0.87 and 10 validated in mice and therefore are not included in our prediction model in B. **[B]** Predictive model for WMMS and phenotypic outcomes of prenatal death and post-natal hydrocephalus for 124 major and minor mutations. The model is based on the following four assumptions. i.WMMS is linearly proportionate to neurological disease (y = 0.76*x - 2.5; r^2^ 0.9, based on findings of Fig. 6B) ii. A lower attenuated phenotypic threshold is 26% prenatal fatality and 31% post-natal hydrocephalus associated with A394V, WMMS -0.87. iii. The upper threshold of WMMS 10 is capped at 90% prenatal fatality, the minimal disease level achieved by G561R,S562F (WMMS >10). iv. Formate supplementation rescues prenatal death caused by severe mutation to restore attenuated phenotype. Based on these assumptions, the model predicts that increase in WMMS of a neurogenic mutation predicts increase in prenatal death enumerated by progressive decrease in progeny yields that can be restored by formate supplementation and which does not mitigate glycine levels. Note: measurable post-natal hydrocephalus may also be observed at days 15-45 post birth for all WMMS, unless pups die before symptoms are detected (such as within 24h, as reported for G561R,S562F).

We next incorporated these WMMS analyses into a model predicting the phenotypic outcome of disease in mice (Fig. 8B). The lower threshold of the model is ∼26% prenatal fatality and 31% post-natal hydrocephalus for WMMS -0.87, as validated by engineering A394V in mice. The upper threshold of WMMS 10 was capped at 90% prenatal fatality since this was the minimal disease level achieved by severe G561R,S562F double mutation, WMMS >10. Formate supplementation rendered the highly severe state back to 29% prenatal fatality and 34% post-natal hydrocephalus (akin to the attenuated state), suggesting that lower levels of prenatal disease induced by mutations with -0.87<WMMS<10 should also be rescued by formate.

In Fig. 8A, 124 NKH mutations show WMMS between -0.87 and 10. Hence, the model in Fig. 8B predicts that all may be successfully expressed in the C57BL/6 mice to present prenatal and postnatal neurological outcomes, with prenatal disease increasing with rising WMMS. For mutations where prenatal disease is high (7.44<WMMS<10), pretreatment of dams with formate supplementation may be needed to yield productive levels of viable progeny. Severe mutations in the range 1.3 >WMMS<7.44 range are expected to show higher levels of prenatal death than those seen in attenuated mice. In this range, 10 mutations with high prevalence and/or observed in homozygous state in humans constitute an important subset of the 81 prevalent severe neurogenic NKH missense mutations (WMMS 1.3-7.4, S Table 12). In summary, Fig. 8B provides a model for large scale prediction of prenatal disease outcomes across hundreds of neurogenic NKH mutations and suggests that high WMMS mutations underlie prenatal disease that is fatal unless rescued by formate.

## Discussion

Our work lays the basis to broadly understand the function of GLDC, a key mitochondrial protein. Multiple new and rapid *in silico* mutation-based tools and mouse models serve assessment and analysis of neurological disease in NKH, as well as the acute need to predict prenatal and postnatal outcomes and develop new, rational treatments for a deadly neurometabolic disorder. In general, in the development of genetic mouse model studies, the dominant criteria for the selection of a mutation is the prevalence of the mutation and whether the residue under consideration is conserved in the mouse protein. Here, WMMS scores enabled a robust comparison between human neurogenic NKH mutations and their mouse counterparts. Unexpectedly, the equivalence of severity could be predicted for only ∼60% of conserved mutations, thereby improving mutant selection for the development of biological models. When the WMMS yielded multiple choices (such as in the selection of a highly severe mutation to be expressed in mice), we depended on paradigms lodged in evolutionary structural biology in conjunction with WMMS. Combining *in silico* analyses of WMMS and human clinical outcomes scores with *in vivo* evaluation of an attenuated mutation in mice, enabled defining (i) WMMS 1.3 as the threshold separating attenuated and severe disease in a linear model of disease and (ii) an attenuated but nonetheless symptomatic score range of -0.87 and 1.3, expected to apply to both mouse and human mutations.

Human NKH is a complex disease with over 50 symptoms that have recently be stratified into four disease domains^17^. Yet the mouse models generated in this study presented one major fatal brain defect of hydrocephalus. Acquired hydrocephalus has been reported in patients^36,37^. However additional disease domains, particularly seizures and cognition, need to be assessed in future studies. It is possible that these may spontaneously emerge upon testing of mutations 1.3<WMMS<7.4, that are expected to be more severe than the attenuated mutation tested here. In this range, R515S (WMMS 3.5), the most prevalent clinically severe mutation and found in homozygous alleles of patients, is of particular interest. As shown in the model in Fig. 8B, WMMS predicts that its mouse counterpart R520S is likely to yield substantial levels of prenatal death compared to attenuated mutations but nonetheless also produces sufficient levels of viable progeny that may show multiple disease domains in addition to the cerebral defect of hydrocephaly. Although R515S is a known severe mutation, it is not a residue in the active site region or other regions that have been previously annotated. Thus, its validation in mice may also reveal information on new functional domains in GLDC.

The genetic background of the mouse can also play a key role in the symptoms expressed in a strain. It is therefore possible that C57BL/6 strain, selected for ease of genetic engineering, may not be the optimal background to capture a broader range of NKH symptoms. Nonetheless, we were able to predict a model relevant to hundreds of mutations, where severe mutations cause high levels of prenatal disease, suggesting they limit a critical step in embryonic development. That the severe mutant was rescued by formate treatment of dams expected to normalize one-carbon metabolism strongly suggests that prenatal death in absence of formate may be due to neural tube defect. While deletion of *Gldc* has been previously shown to cause neural tube defect that may be rescued by formate treatment of dams^35,38^, we provide the first evidence that severe mutation-induced neurological disease in NKH is principally due to prenatal defects (milder forms of which could induce post-natal hydrocephalus) and that the WMMS in the C57BL/6 strain provides a predictor of fatal, prenatal defect/disease.

A direct consequence of defect in GLDC is the failure to cleave glycine. Glycine elevation is seen in the blood of NKH patients. But plasma glycine increases induced by the attenuated mutation and formate-rescued severe double mutation, were comparable as detected between 30-60 days. Age-dependent glycine was detected in more mature animals, but additional studies are required to determine whether the elevation of glycine is associated with other symptomatic domains (such as seizures) not yet assessed in our work in mice. Nonetheless, our studies establish that while glycine is an age-dependent signifier of GLDC defect, treatment for normalization of single carbon metabolic deficiency (a second expected consequence of *Gldc* mutation), diminishes prenatal defects that underlie mutation-based, neurologic disease. This suggests that the observed clinical benefit of sodium benzoate and dextromethorphan that respectively reduce glycine or antagonize its action, could well be due to off-target effects. Alternatively, it is possible that in the mouse models, seizure is not a consequence of prenatal disease. Studies with additional NKH mutations and possibly other mouse genetic backgrounds are needed to further solve the complex neurometabolic basis of NKH. Our work provides a new framework for large scale understanding of mutation functions and the prediction that severity of a neurogenic mutation is a direct measure of pre-natal disease in NKH mouse models.

## Acknowledgments

We thank Mehdi Ghorbal for advice on CRSPR-Cas 9 and Dr. Abhishek Trivedi and Ashley Van Avermaete for assistance with genotyping and mouse colonies. We thank the Freimann Life Sciences Center, University of Notre Dame for breeding and maintaining mouse colonies. We thank Taconic Biosciences and The Jackson Laboratory for engineering transgenic mice and providing founder mice and the Wistar Institute Proteomics and Metabolomics, Philadelphia, USA for conducting glycine measurements. The Thermo Q-Exactive HF-X mass spectrometer used by the Wistar Institute was purchased with NIH grant S10 OD023586. We thank Stefan Freed and Dr. Shaun Lee and all members of the Haldar lab for helpful advice and discussion. We thank Dr. Kalipada Pahan for preliminary assistance with glycine measurements.

## Author Contribution

JF – conceptualization, curation and analysis of computational and mouse data; designed, executed and analyzed all computational methods, software writing and *in silico* data curation and prediction; visualization of results; drafting and editing of manuscript.

SA –conceptualization, curation and analysis of mouse data; supervision, genotyping strategy and assay development; organ harvest; glycine analysis; breeding and management of mouse colonies; visualization of results, drafting and editing of manuscript.

AM – mouse genotyping and observation; colony management; organ harvest; data visualization.

KH – overall conceptualization and project supervision, design and development of mouse and computational studies; formal analysis of all predictive *in sili*co and *in vivo* mouse data, visualization of results; drafting and editing of manuscript, funding acquisition.

All authors read the manuscript and commented on it.

